# Therapeutic Potential of Flavonoids and Rutin Against Inflammation on a Huntington Animal Model

**DOI:** 10.64898/2025.12.02.691965

**Authors:** David Calderón Guzmán, Norma Osnaya Brizuela, Maribel Ortiz Herrera, Hugo Juárez Olguín, Armando Valenzuela Peraza, Alberto Rojas Ochoa, Ernestina Hernández García, Rafael Coria Jiménez, Daniel Santamaria del Angel

**Affiliations:** Laboratorio de Neurociencias, Instituto Nacional de Pediatría (INP), Mexico City, México; Laboratorio de Bacteriología Experimental. INP, Mexico City, México; Departamento de Farmacología, Facultad de Medicina, Universidad Nacional Autónoma de Mexico, Mexico City, Mexico; Laboratorio de Oncología Experimental INP, Mexico City, México; Laboratorio de Farmacología, INP, Mexico City, Mexico

**Keywords:** Brain, Huntington animal model, Oxidative stress, inflammation, Biogenic amines

## Abstract

This study tested the hypothesis that flavonoids (diosmine, hesperidin and rutin®) play a protective role in the brain and duodenum against 3-NPÁs free radical induced inflammation. This was achieved by measuring the levels of GABA, 5-HIAA, dopamine and some inflammation and oxidative stress markers. Male young Wistar rats (weight 60g) received *Salmonella thyphimurium* ATCC14028 1×10^6^ CFU/g every week, for two consecutive weeks, plus the following treatments: group A, 9% NaCl 0. (Control); group B, diomine (300mg) + hesperidin (33.3g) + rutin (150mg/kg); group C, 3-NPA; group D, 3-NPA + mix of flavonoids with 1ml of rutin® per rat. 3-NPA administration was at 24mg/kgW by intraperitoneal route and the flavonoids were orally administered at every 48 hours for 15 days. At the moment of euthanasia, the blood was obtained to assess Interleukine-6, glucose, triglycerides and hemoglobin levels. Brain and duodenum were obtained to measure GABA, dopamine, 5-HIAA, lipoperoxidation, reduced glutathione (GSH), total ATPase concentrations and catalase activity using validated methods. The brains, stomach and duodenum were dissected for histological analysis. In group A, there is a decrease in interleukine-6 levels (p=0.009). In the cortex region of animals in group C, dopamine experienced a significant decrease (p=0.012). GABA increased (p=0.001) in cerebellum regions of animals in the groups A, B and C. ATPase activity increased (p = 0.013) in cerebellum regions in animals of groups C and D. Lipoperoxidation diminished (p=0.043) in cerebellum region of group D. Catalase activity diminished (p=0.043) in Cortex region of animals in groups B and D, and (p=0.002) in cerebellum region of animals in groups A and B. Besides, histological changes revealed marked lesions of neuronal cells in experimental animals treated with nitro propionic acid. The protective role of flavonoids and rutin compounds on inhibition of the inflammatory response and correction of the fundamental oxidant/antioxidant imbalance in animals suffering from Huntington diseases are important vistas for further research.

## INTRODUCTION

Neuro diseases propose excessive stimulation of excitatory neurotransmitters that brings about augmentation of the concentration of intracellular calcium ions. This situation triggers lethal pathway for reactive oxygen (ROS) and nitrogen species production [1], provoking oxidative stress and release of nitric oxide (NO^ˉˉ^) species. NO^ˉˉ^ not only causes physical changes in the structure of the mitochondria and negatively impact on their functions, but also leads to DNA damage [2]. 3-nitroptopionic acid (3-NPA) is a potent mitochondrial toxin and generally causes the damage of mitochondria and DNA. In Wistar rats, 3-NPA promotes the degeneration of neurons. Its systemic delivery in rats heralds Huntington disease model [3], where it works in the basal ganglia using up the store of monoamine neurotransmitters, such as serotonin, norepinephrine, and dopamine [4]. Apart from its toxic potential, NO^ˉˉ^, in the low nanomolar range of 10^ˉ8^ to 10^ˉ6^ M could function as a neuromodulator, however any deviation from this amount may trigger fatal consequences on the cells through the formation of nitroso-glutathione (NOGSH) and increase of in oxidative stress [5]. The fact that cellular structural components are susceptible to damage by free radicals [6], more specially the membrane lipids [7], the central nervous system (CNS) must be protected from NO^ˉˉ^ imbalance through the maintenance of the right amount of antioxidants. This point is extremely important during development, because this is the time brain metabolism and growth is at peak level [8]. Therefore, it is necessary to regulate energy and glucose homeostasis via the maintenance of dopaminergic system in hypothalamic neurocircuits and higher brain circuits [9]. Apoptosis of macrophages in the body is found during systemic infection, and the activation of kinase pathways leads to balanced pro- and antiapoptotic regulatory factors in the cell. In the intestine, *Salmonella* mediates macrophagic death by caspase-1 activation, which also releases interleukins. The above situation bolsters inflammation and triggers chemiotactic influx of phagocytic cells such as macrophages, thus carrying the infectious agent to tissues outside the intestine [10]. However, this disseminative channel is not the only pathway Salmonella can spread to systemic level. Some strains can invade intestinal epithelial cells through the “zipper” and “trigger” pathways leading to seriously fatal infections and production of extensive cytopathology that reaches out to many systemic organs [11]. In any case, the overall effectors involved in this dissemination pathways still remain poorly clear.

In the context of neurodegenerative diseases, phytonutrient compounds, such as flavonoids, are natural products with preventive potentials. These diseases share the same pattern based on progressive deterioration of dopaminergic (DA) neurons [12]. The flavonoids activate the antiapoptotic pathways, such as P13K/AKT and NF-_K_B, that are critical intracellular pathways geared towards the regulation of cell survival, growth and proliferation, as well as the protection of the mitochondria from dysfunction and activation of neurotrophic factors. Findings of Burda and Oleszek [13], suggest that flavonoids with hydroxyl group in position C-3 possess elevated antioxidant activity, while the hydroxyl in C-4 possesses anti-free radical activity, and that these compounds have shown antimicrobial activity [14]. The flavonoids as diosmine (figure 1) and hesperidin (figure 2) are natural polyphenolic compounds with antioxidant and anti-inflammatory effects [15]. The intake of Rutin® (figure 3) is recommended for its neuroprotective roles helping to downplay neurodegenerative disorders due to its antioxidant activities [16]. Particularly, rutin® has some protective effects in Huntingtońs Disease models [17] although the underlying mechanisms are still unknown.

**Figure 1.**
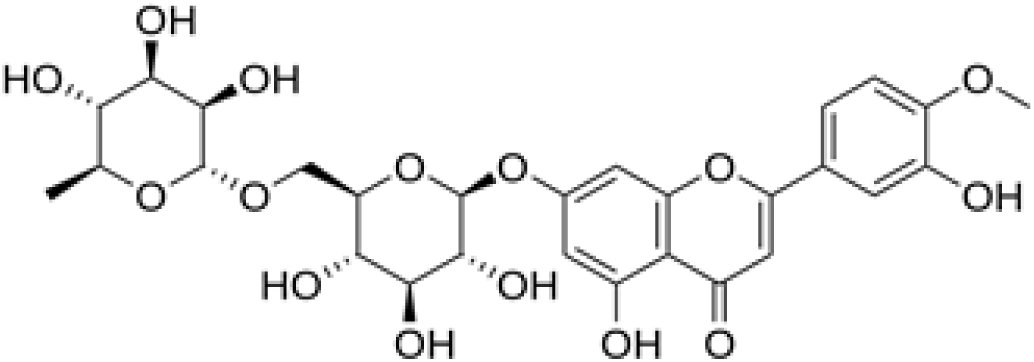
Diosmine. 3’,5,7-Trihydroxy-4’-methoxyflavone 7-rutinoside.

**Figure 2.**
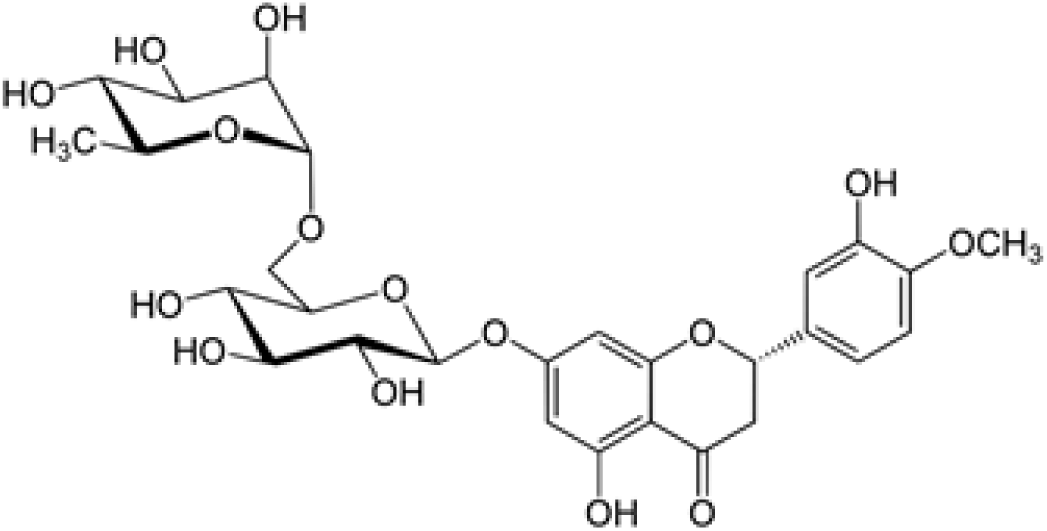
Hesperidine. (2*S*)-5-hydroxy-2-(3-hydroxy-4-methoxyphenyl)-7-[(2*S*,3*R*,4*S*,5*S*,6*R*)-3,4,5-trihydroxy-6-{[(2*R*,3*R*,4*R*,5*R*,6*S*)-3,4,5-trihydroxy-6-methyloxan-2-yl]oxymethyl} oxan-2-yl]oxy-2,3-dihydrochromen-4-one.

**Figure 3.**
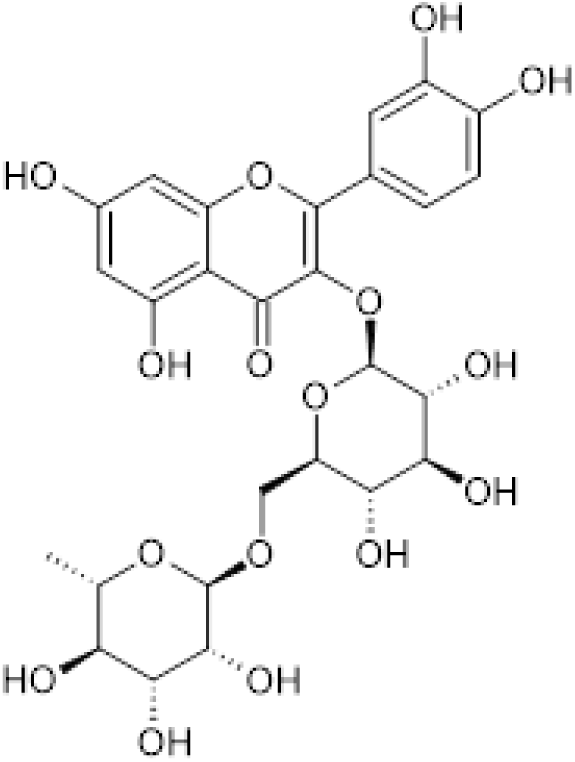
Rutin. [3’,4’,5,7-Tetrahydroxy-3-(α-L-rhamnopyranosyl-(1→6)-β-D-glucopyranosyloxy]

Plasma membrane phospholipids in brain are in close contact with structural proteins that are embedded in the lipid bilayer [18], from which Na^+^, K^+^ ATPase is responsible of keeping the ionic interchange through this bilayer by the stimulation of Na^+^ and K^+^ flows [19]. The inhibition of the Na^+^, K^+^ ATPase activity induces excitatory amino acid release within the Central Nervous System (CNS) [20].

Taken the above reports and findings as a background, the purpose of the present study is to compare the protective effect of flavonoids (diosmine and hesperidin) in combination with rutin® on the levels of dopamine, GABA, 5-HIAA and on oxidative stress and antiinflammatory markers in brain regions, duodenum and stomach of animal model with experimentally induced inflammation and Huntingtońs disease.

## MATERIAL AND METHODS

### Experimental animals

Animals were purchased from certified bioterium of Instituto Politecnico Nacional, Mexico City. The animals were placed in four meshed plastic cages, each containing eight rats and were exposed to 12 h light-dark cycle and natural environmental conditions. Free access to pelleted laboratory rodent feed (Purine 5001) and water was allowed during the experiment. Before the study, the animals were allowed 1 - 2-week period of acclimatization to the animal house facility conditions with food and water. Animal management and care were conducted according to the National and International guidelines of animal care. This study protocol was approved with the reference number 026/2022.

### Chemicals

Thiobarbituric, Glutathione, catalase, ATP, GABA, Dopamine, 5-HIAA and Ortho Pthaldialdehyde were acquired from Sigma-Aldrich, St. Louis, MO, USA. Hydrochloride acid, Sulfuric acid, Nitric acid, Bisulfite, Trichloro acetic acid, Sodium phosphate, Magnesium chloride and Methanol were purchased from Merck, Darmstad, Germany. Triglycerides and glucose Roche devices were used in the study. Catalase Assay Kit was from Cayman Chemical Company, and Rat IL6 and Interleukin-6 Elisa Kit were obtained from OriGene Technologies Inc.

### Experimental model

Thirty young male Wistar rats (60 g) were separated into 4 groups and treated as follows: Group A, 0.9% NaCl + *Salmonella T* (control). Group B, *S. typhimurium* + Diosmine/hesperidine/Rutin® (1 ml); group C, *S. typhimurium* + 3-NPA (24 mg/kg); group D, *S. typhimurium* + Diosmine/hesperidine/Rutin (1 ml) + 3-NPA (24mg/kg) per rat. Diosmine/hesperidine/Rutin® was orally administered every 48 hours for 15 days. Live culture of *Salmonella typhimurium (S. typhimurium)* ATCC14028, 1 x 10^6^ colony-forming units/rat (CFU/rat) was given once a week in two doses, and 3-NPA in a single dose at the end (Experimental design). 120 minutes after receiving the drugs, the animals were put under anaesthesia (sodium pentobarbital 50 mg/kg) and euthanized with guillotine to obtain the brain, stomach and duodenum, and then put in saline (NaCl 0.9 %) at 4 °C. Eight animals (two rats for each group) were stained with hematoxylin-eosin to evaluate the histological abnormalities. The blood was assessed to measure Interleukin-6, triglycerides, hemoglobin and glucose. Brain was dissected into cortex, hemispheres and cerebellum. Brain regions, stomach and duodenum were put in 5 volumes of 0.05 M TRISHCl, pH 7.4 to evaluate lipoperoxidation (TBARS), total ATPase and catalase. An aliquot was homogenized in 0.1 M perchloric acid (HClO_4_) (50:50 v / v) to evaluate γAminobutyric acid (GABA), reduced glutathione (GSH), dopamine and 5-hydroxyindole acetic acid (5-HIAA) concentrations.

### Experimental design and treatment

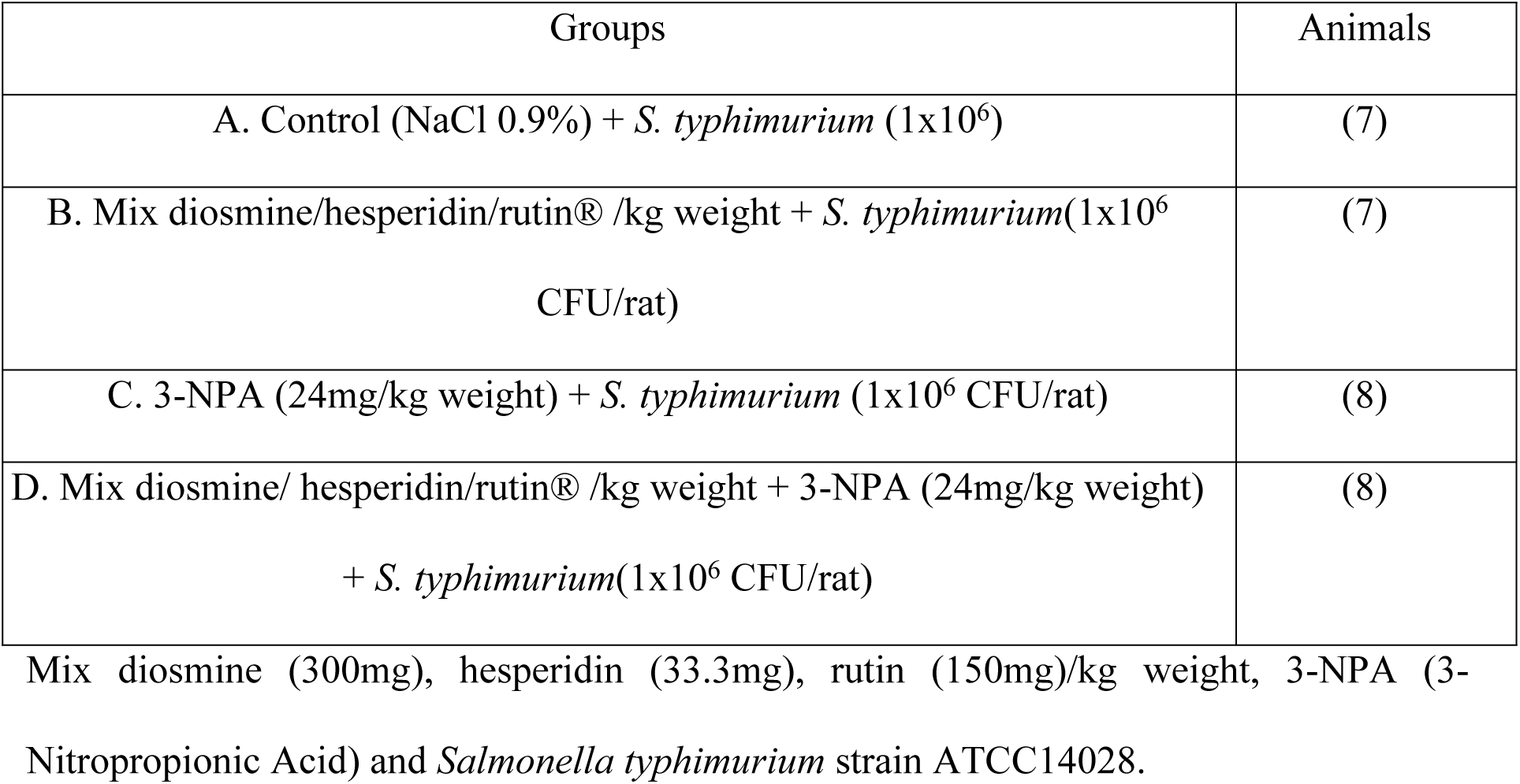

| Groups | Animals |
| --- | --- |
| A. Control (NaCl 0.9%) + *S. typhimurium* (1x10<sup>6</sup>) | (7) |
| B. Mix diosmine/hesperidin/rutin® /kg weight + *S. typhimurium*(1x10<sup>6</sup> CFU/rat) | (7) |
| C. 3-NPA (24mg/kg weight) + *S. typhimurium* (1x10<sup>6</sup> CFU/rat) | (8) |
| D. Mix diosmine/ hesperidin/rutin® /kg weight + 3-NPA (24mg/kg weight) + *S. typhimurium*(1x10<sup>6</sup> CFU/rat) | (8) |

Mix diosmine (300mg), hesperidin (33.3mg), rutin (150mg)/kg weight, 3-NPA (3-Nitropropionic Acid) and *Salmonella typhimurium* strain ATCC14028.

### Inoculation of rats with Salmonella thyphimurium strain ATCC14028

The *S. thyphimurium* inoculated in the rats were obtained from strain bank (ceparium) of Experimental Bacteriology laboratory of National Institute of Pediatrics, Mexico City. Following its re-identification, an aliquot was taken from a maintenance medium and injected in a culture medium of Salmonella Shigella (SS agar). This was subjected to 18- to 24-hour incubation at an ambient temperature of 37 °C using bacteria incubator, Zhengzhou Nanbei instruments, Henan, China. Subsequently, proven colonies containing *S. thyphimurium*, after reconfirmation through conventional biochemical tests, were selected and inoculated in TSA (Trypticasein Soya Agar). This was subjected to 18-hour incubation at 37 °C. Using Hyssop, the collection of bacterial biomasses was made. The masses were suspended in PBS buffer, pH = 6.8. Subsequently, with DU 640 spectrometer (Beckman, USA) the masses were adjusted to an AS_450nm_= 0.175 (equivalent to 3 x 10^8^ UFC / ml) and diluted to a concentration of 1 x 10^6^ UFC / ml [21]. A non-lethal volume (1 ml) was taken and administered to each animal using orogastric tube.

### Technique to measure glucose and triglycerides in blood

Glucose and triglycerides were measured using 20 µl of tail-end blood, twice collected after the treatment. The collection was made without anticoagulant. This volume of blood was smeared on a reactive filter paper in Accu-Chek active (Roche Mannheim Germany) equipment. The glucose and triglyceride concentrations were read and reported in mg/dL.

### Measurements of Interleukin (IL-6)

The measurement of IL-6 was carried out at the end of treatment. 3 mL of fresh blood was obtained from the heart by cardiac punction after anesthesia. This was centrifuged at 3,500 rpm for 10 min in a clinical centrifuge (HERMLE Labnet, Z 326 K). The plasma obtained was processed with Rat IL6/Interleukin-6 Elisa Kit from OriGene Technologies Inc. The samples were read in triplicate at 450 nm in a microplate reader. Molecular devices Spectra max plus 384 and software SoftMax Pro 6.0 were used and the concentration was expressed in pg/ml.

### Dopamine concentration determination

The determination of dopamine (DA) concentrations was carried out with HCLO_4_ homoginzed tissue supermatant resulting from a 10-minute centrifugation at 9,00 rpm performed with a microcentrifuge (Hettich Zentrifugen, model Mikro 12-42, Germany). The technique employed was reported by Calderon et al, [22]. A portion of the HClO_4_ supernatant was mixed with 1.9 ml volume of chemical buffer solution (0.003M octylsulphate, 0.035 M KH_2_PO_4_, 0.03 M citric acid, 0.001 M ascorbic acid) in a test tube. In total darkness at room temperature, the incubation of the mixture was performed for a 5minute period. Thereafter, the reading of the samples was carried out using a spectrofluorometer (Perkin Elmer LS 55, England) with an excitation wavelength of 282 nanometers and electromagnetic radiation at a specific wavelength of 315 nanometers The FL Win Lab version 4.00.02 software was used. Using a previously standardize curve, the concentration values of DA was deduced and this was reported in nanomol per gram (nMoles/g) wet tissue.

### Measurement of γ-Aminobutyric acid

The determination of the concentration of γ-Aminobutyric acid (GABA) was carried out with the HCLO_4_ homogenized tissue supernatant previously subjected to a 10-minute centrifugation at a revolution of 9,000 per minute using a microcentrifuge (Hettich Zentrifugen, model Mikro 12-42, Germany). The technique employed in this determination was developed by Hsieh et al, [23]. An aliquot of the HClO_4_ supernatant and work solution (Buffer of Boric acid 0.1M pH 9.3 + MeOH + Orto-Phthalaldehyde + Mercaptoethanol) was collected and loaded in a test tube. Subsequently, this was subjected to 5-minute incubation at room temperature in total darkness. Following the incubation, GABA concentrations in the samples were fluorometrically read using FL Win Lab version 4.00.02 of Perkin Elmer LS 55 (England) spectrofluorometer with an excitation wavelength of 340 nanometers and electromagnetic radiation at wavelength of 455 nanometers. Using a previously standardize curve, the concentration values of GABA was deduced and this was reported in nanomole per gram (nMoles/g) wet tissue.

### 5-HIAA concentration Determination

The levels of 5-HIAA were evaluated using the floating tissues of the brain regions previously mixed with HClO_4_ (2:1 v/v). The tissues were made to undergo a 10-minute centrifugation at a revolution of 10,000 per minute using a micro centrifuge (Hettich Zentrifugen, model Mikro 12-42, Germany). To process the brain regions, an aliquot of each region was fed to Perkin Elmer LS 55 fluorometer that processed the tissues at an excitation wavelength of 296 nanometers and an emission of 333 nanometers and FL Win Lab version 4.00.02 software was to determine the concentration values [24]. The values were extrapolated in a standard curve previously standardized and reported in nM/g of wet tissue.

### Reduced Glutathione (GSH) concentration Determination

To measure the concentration of GSH, the HCLO_4_ homogenized tissue supernatant that was previously centrifuged with MIkro 12-42 centrifuge (Germany) during a 5-minute period at a revolution of 9,000 per minute was used applying Hissin and Hilf modified method [25]. In a phosphate buffer [pH 8.0, EDTA 0.2%] with a capacity of 1.8 mL, the supernatant aliquot (20 μL) was mixed with ortho-phthaldehyde (100 mL) and methanol (1 mg/mL). The mixture was put in a test tube and subjected to a 15-minute incubation at room temperature in absolute darkness. Perkin Elmer LS 55, software FL Win Lab 4.00.02 version, that possesses an excitation wavelength of 296 nanometers and an emission of 333 nanometers was used to read GSH concentration in the centrifuged samples. From a previously standardised curve the concentration of GSH was extrapolated and reported in nanomole per gram (nMoles/g) wet tissue.

### Total ATPase Measurement

Adenosine triphosphate enzyme activity was analyzed based on Calderón and colleagueś method [26]. The analysis was conducted with 1 mg (10%) weight-by-volume (w/v) of the tissues of brain, duodenum and stomach that were previously homogenized in tris-HCl 0.05 M pH 7.4. The 1 mg of each of the tissues were put in a solution containing 3 mM MgCl_2_, 7 mM KCl, and 100 mM NaCl and subjected to a 15-minute incubation. Following this incubation, 4 mM tris-ATP was added to the solution. With Dubnoff Labconco shaking water bath, a second-round 30-minute incubation of the new solution was carried out at 37 °C. To stop the reaction, 100 µL (10%) weight-by-volume (w/v) of trichloroacetic acid was used. Subsequently, the samples were centrifuged at 100 g for 5 minutes at 4 °C. The measurement of inorganic phosphate (Pi), a product of ATP hydrolysis by ATPase, was carried out in triplicates using one supernatant aliquot as proposed by Fiske and Subarrow [27]. The supernatant absorbance reading was made with BECKMAN DU 640 spectrophotometer at 660 nanometers. The absorbance was expressed as mM Pi/g wet tissue per minute.

### Measurement of Catalase

The determination of catalase was made with catalase kit (Cayman Chemical®) using the modified technique of Sinha [28]. Each brain region (cortex, hemispheres, cerebellum/medulla oblongata), stomach and intestine were homogenized in 3 mL of trisHCl 0.05 M pH 7.4 buffers. From the diluted homogenates, 100 µL was taken. The samples were read in triplicate at 570 nm in a microplate reader. Molecular devices Spectra max plus 384 and software SoftMax Pro 6.0 were used. Catalase activity was expressed in µM/g of wet tissue.

### Lipid peroxidation (TBARS) determination technique

The measurement of TBARS was made using the brain tissues homogenized in tris-HCl 0.05 M pH 7.4 buffer solution in accordance with the modified technique of Gutteridge and Halliwell [7]. 1mL of the homogenized brain tissue sample was mixed with 2mL solution containing 1.25 g of thiobarbaturic acid (TBA), trichloroacetic acid (TCA) 40 g, and concentrated hydrochloric acid 6.25 mL that was diluted with deionized water 250 mL. The mixture was put in a Thermomix 1420 and subjected to 30-minute heat. Thereafter, it was cooled for 5 minutes in an ice bath followed by 15-minute centrifugation at 700 gforce (Sorvall RC-5B Dupont). Subsequently, the reading of the floating tissue absorbance was carried out in triplicate at 532 nanometer using BECKMAN DU 640 spectrophotometer. The reactive substance concentration to TBARS was reported as µM of Malondialdehyde/g of wet tissue.

### Histological analysis in brain regions, stomach and duodenum

The brain, stomach and duodenal tissues were gentle cleansed in saline solution (0.9% NaCl) following their extraction with the objective of removing all the sticking debris on them. They were immediately subjected to histological analysis consisting of fixing them for 24 hours in a 10% NBF (neutral buffered formalin) solution. This was followed by washing and removing excess parts on the tissue samples and dehydrating them in a graded series of alcohol. Thereafter, the samples were immersed in xylene to remove the water and alcohol, and then paraffin-embedded. Subsequently, they were cut to a thickness of 4-6 millimeters and H&E stained (Hematoxyn and Eosin). Next, the H&E-stained tissues were viewed under stereology microscope (Olympus BX51). Software Stereo Investigator 11 (SI 11), was used to quantify the biochemical indicators [29].

### Statistical analysis

Descriptive statistic tables containing central measures of tendency and dispersion were employed to show the data. Inference analysis was performed to compare the biochemical indicators of the control group with the experimental animal groups. Contrast of hypothesis test such as Fisher analysis of variance (Anova) or Kruskal-Wallis test after variance homogeneity verification was used for this purpose. Tukey-Kramer or Steel-Dwass tests was employed as *Post hoc* contrasts test. Any associated probability value p <0.05 was considered statistically significant. Analysis was performed using Sigma Plot Statistical v12 software [30].

## RESULTS

The result of Interleukin-6, triglycerides, glucose and hemoglobin levels in the blood of rats treated with mix flavonoids + *Salmonella T*. in the presence of 3-nitropropionic acid are presented in Table 1. In animal groups treated with Mix flavonoids + *Salmonella typhimurium* + 3 -NPA, Interleukin - 6 levels decreased (p = 0.009) with significant differences when compared with the control group.

**Table 1.**
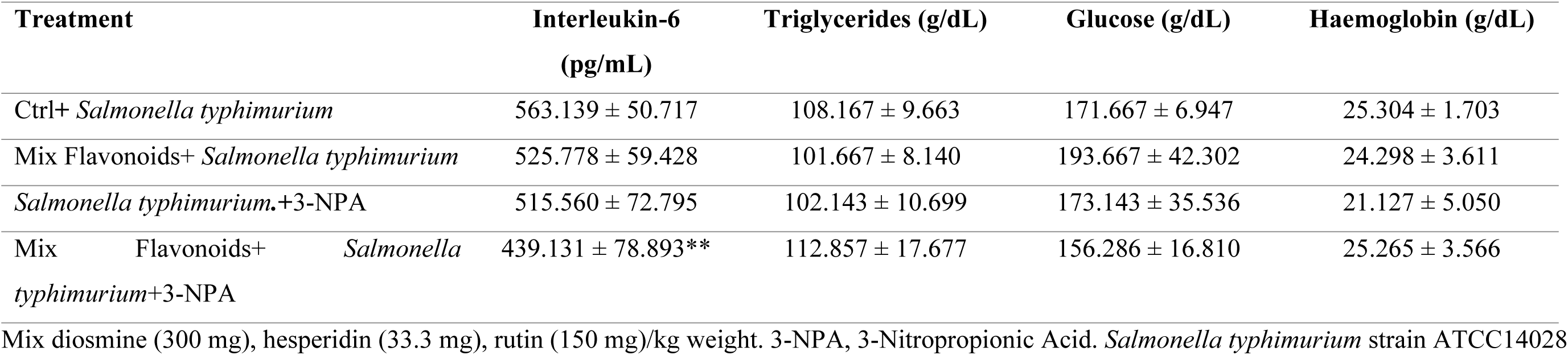
Blood levels of Interleukin-6 in rats with *Salmonella thyphimurium*: Mix Flavonoids+ *Salmonella typhimurium.*+3-NPA vs Ctrl+ *Salmonella typhimurium* **p=0.009. Triglycerides, Glucose and Hemoglobin p=N.S.

Dopamine levels in brain regions of rats treated with mix flavonoids + *Salmonella typhimurium* in the presence of 3-nitropropionic acid are presented in Table 2. Dopamine diminished significantly (p=0.012) in cortex region of animals that received *Salmonella typhimurium* combined with 3-nitropropionic acid when compared with the control group. GABA levels in brain regions of animals treated with mix flavonoids + *Salmonella typhimurium* in the presence of 3-nitropropionic acid are shown in Table 3. GABA increased significantly (p=0.001) in Cerebellum regions of animals that received *Salmonella typhimurium* alone, or combined with Mix Flavonoids, or 3-NPA compounds when compared with the combination of mix Flavonoids+ *Salmonella typhimurium* + 3NPA group.

**Table 2.**
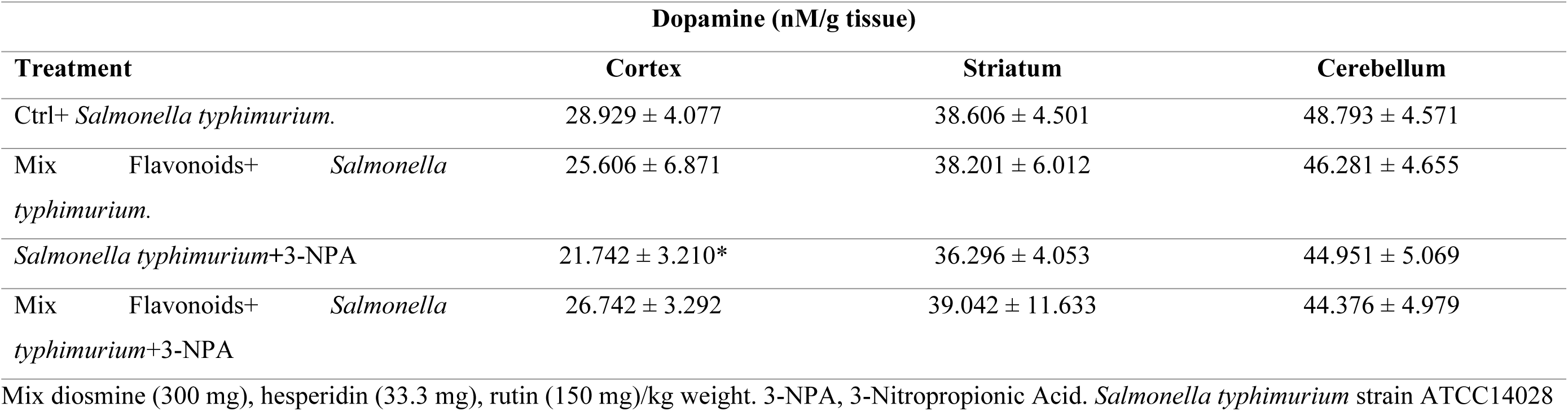
Brain levels of Dopamine in rats with *Salmonella thyphimurium*: Cortex: *Salmonella typhimurium* + 3 - NPA vs Ctrl+ *Salmonella typhimurium* *p = 0.012. Striatum and Cerebellum p = N.S.

**Table 3.**
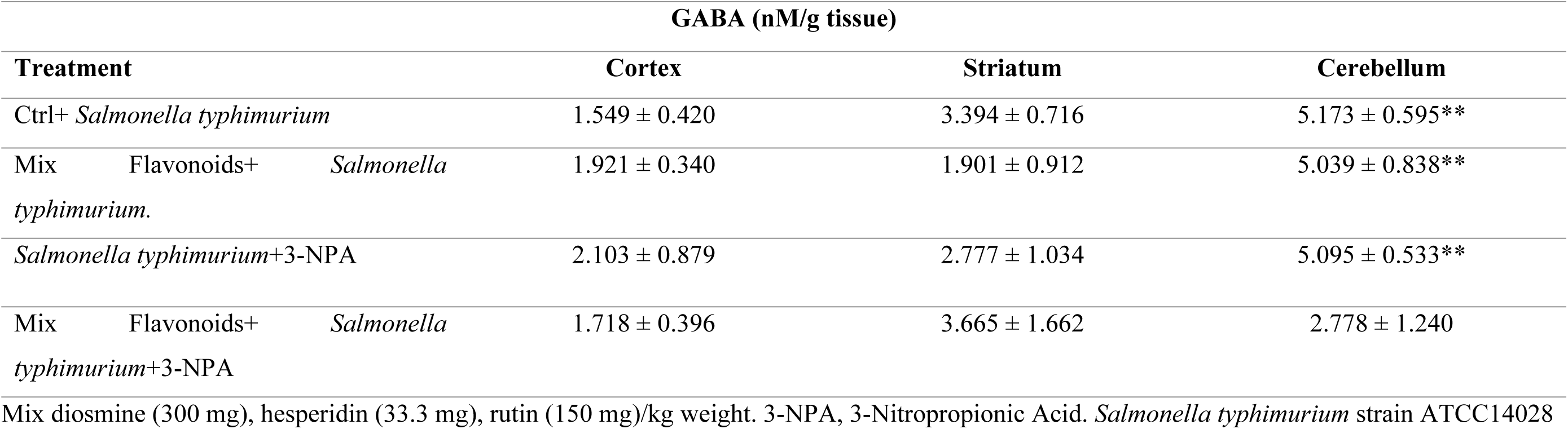
Brain levels of GABA in rats with *Salmonella thyphimurium*: Cerebellum: Ctrl+ *Salmonella typhimurium,* Mix Flavonoids+ *Salmonella typhimurium.* and *Salmonella typhimurium* + 3-NPA vs Mix Flavonoids + *Salmonella typhimurium* + 3-NPA **p=0.001. Cortex and Striatum p = N.S.

Table 4 shows the levels of 5-HIAA in brain regions of rats treated with mix flavonoids + *Salmonella typhimurium* in the presence of 3-nitropropionic acid, where no significant difference (p > 0.05) was observed between them and the control group.

**Table 4.** Brain levels of 5-HIAA in rats with *Salmonella thyphimurium*: Cortex, Striatum and Cerebellum p=N.S.

| 5-HIAA (mM/g tissue) |  |  |  |
| --- | --- | --- | --- |
| Treatment | Cortex | Striatum | Cerebellum |
| Ctrl+ *Salmonella typhimurium*. | 1.356 ± 0.299 | 1.816 ± 0.380 | 1.759 ± 0.223 |
| Mix Flavonoids+ *Salmonella typhimurium* | 1.299 ± 0.099 | 1.738 ± 0.232 | 1.875 ± 0.134 |
| *Salmonella typhimurium*+3-NPA | 1.099 ± 0.142 | 1.819 ± 0.312 | 1.854 ± 0.228 |
| Mix Flavonoids+ *Salmonella typhimurium* +3-NPA | 1.264 ± 0.200 | 1.909 ± 0.144 | 1.869 ± 0.199 |

Mix diosmine (300 mg), hesperidin (33.3 mg), rutin (150 mg)/kg weight. 3-NPA, 3-Nitropropionic Acid. *Salmonella typhimurium* strain ATCC14028

GSH levels in brain regions and duodenum of rats treated with mix flavonoids + *Salmonella typhimurium* in the presence of 3-nitropropionic acid are presented in Table 5. There were no significant differences (p > 0.05) between the levels in the experimental animals and in the control group. However, GSH levels was observed to decreased only in duodenum.

**Table 5.** Brain levels of GSH in rats with *Salmonella thyphimurium*: Cortex, Striatum, Cerebellum and Duodenum p=N.S.

| GSH (nM/g tissue) |  |  |  |  |
| --- | --- | --- | --- | --- |
| Treatment | Cortex | Striatum | Cerebellum | Duodenum |
| Ctrl+ *Salmonella typhimurium* | 3.257 ± 0.508 | 4.518 ± 0.564 | 3.327 ± 0.307 | 6.140 ± 3.230 |
| Mix Flavonoids+ *Salmonella typhimurium*. | 2.732 ± 0.500 | 4.285 ± 0.588 | 2.999 ± 0.332 | 5.171 ± 1.108 |
| *Salmonella typhimurium* +3-NPA | 2.825 ± 0.257 | 4.158 ± 0.412 | 3.054 ± 0.433 | 4.931 ± 1.064 |
| Mix Flavonoids+ *Salmonella typhimurium* +3-NPA | 2.953 ± 0.435 | 3.984 ± 0.438 | 3.068 ± 0.490 | 4.787 ± 0.939 |

Mix diosmine (300 mg), hesperidin (33.3 mg), rutin (150 mg)/kg weight. 3-NPA, 3-Nitropropionic Acid. *Salmonella typhimurium* strain ATCC14028

The total ATPase levels (Table 6) in brain regions, duodenum and stomach of rats treated with mix flavonoids + *Salmonella typhimurium* in the presence of 3-nitropropionic acid. ATPase increased significantly (p = 0.013) in cerebellum regions, in animal groups treated with *Salmonella T*. alone or combined with Mix flavonoids or 3-nitropropionic acid with respect to the control group.

**Table 6.**
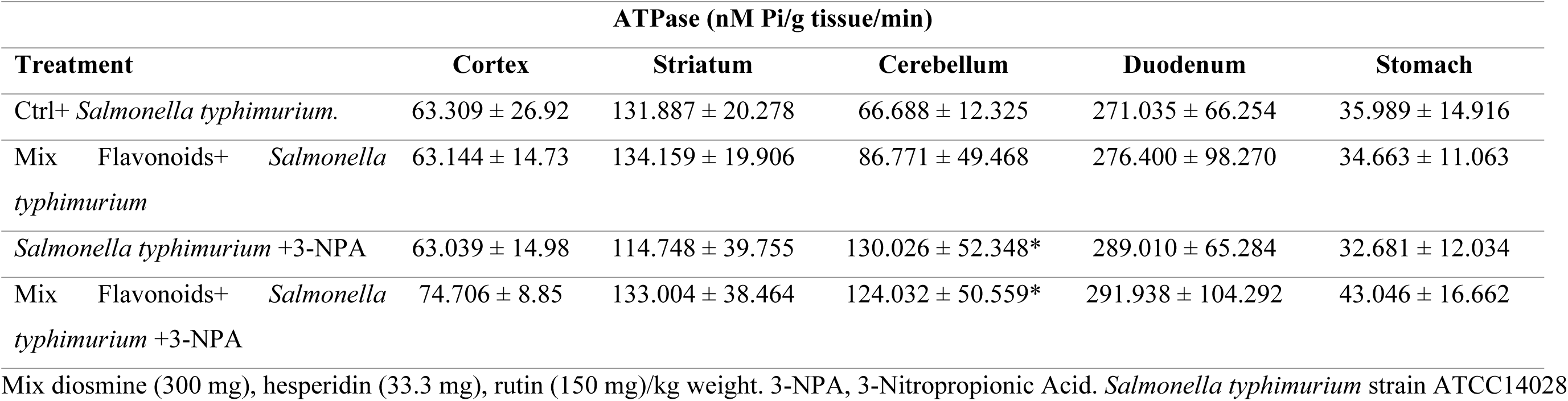
ATPase activity in Brain, Duodenum and Stomach levels of rats with *Salmonella thyphimurium*: Cerebellum: *Salmonella typhimurium* +3-NPA and Mix Flavonoids + *Salmonella T.* + 3-NPA vs Ctrl + *Salmonella typhimurium* *p=0.013. Cortex, Striatum, Duodenum and Stomach p=N.S.

Lipoperoxidation levels in brain regions of rats treated with mix flavonoids + *Salmonella typhimurium* in the presence of 3-nitropropionic acid are shown in Table 7.

**Table 7.**
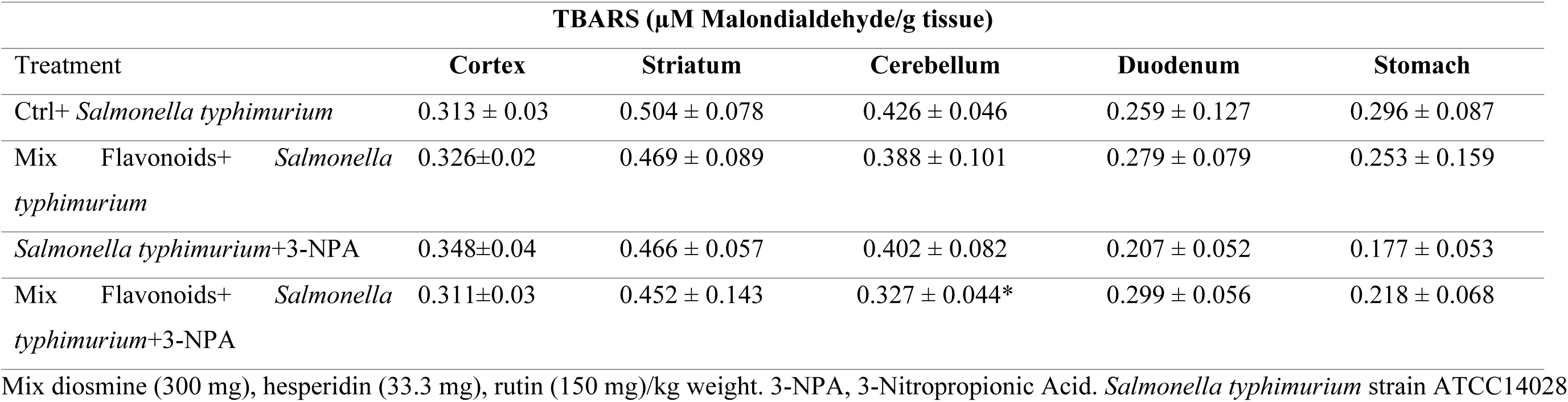
Lipid Peroxidation (TBARS) in Brain, Duodenum and Stomach of rats with *Salmonella thyphimurium* Cerebellum: Mix Flavonoids+ *Salmonella typhimurium* +3-NPA vs Ctrl+ *Salmonella typhimurium* *p=0.043. Cortex, Striatum, Duodenum and Stomach p=N.S.

Lipoperoxidation diminished significantly (p=0.043) in cerebellum region of animals that received *Salmonella typhimurium* combined with mix flavonoids and 3-nitropropionic acid when compared with the control group.

Table 8 depicts the levels of Catalase in brain regions of rats treated with mix flavonoids + *Salmonella typhimurium* in the presence of 3-nitropropionic acid. Catalase activity diminished significantly (p=0.043) in cortex region of animals treated with *Salmonella typhimurium* in combination with mix flavonoids or 3-NPA groups and (p=0.002) in cerebellum region of animals that received *Salmonella typhimurium* alone or combined with mix flavonoids when compared with the same group with 3-nitropropionic acid. Besides, histological changes revealed marked lesions of neuronal cells in experimental animals treated with nitropropionic acid.

**Table 8.**
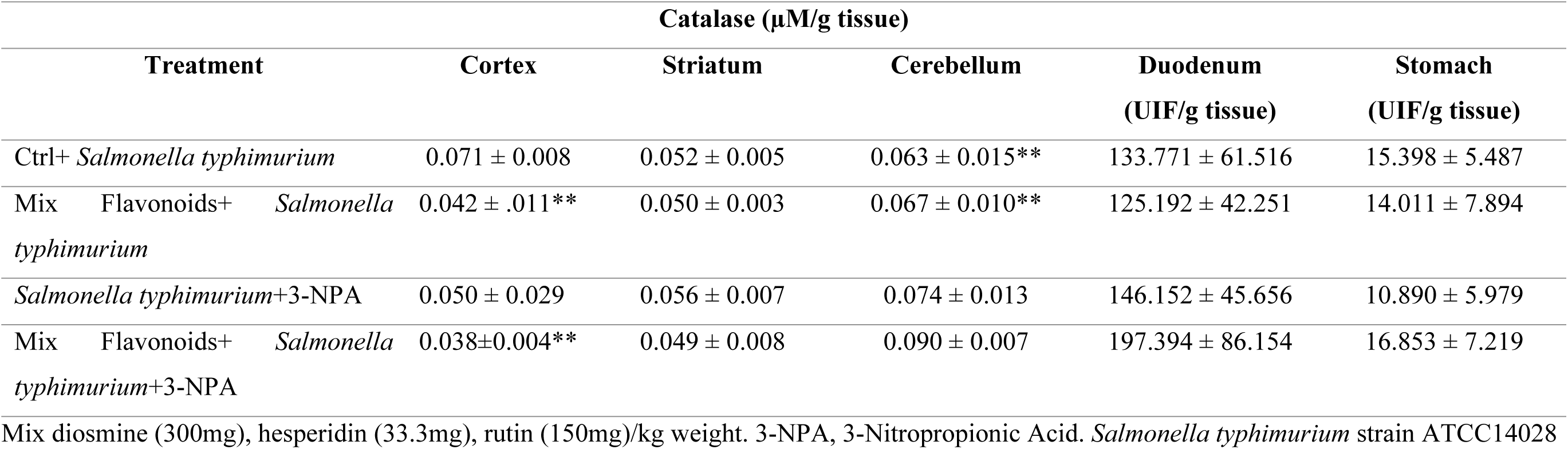
Levels of Catalase in Brain, Duodenum and Stomach of rats with *Salmonella thyphimurium* Cortex: Mix Flavonoids+ *Salmonella typhimurium.* and Mix Flavonoids+ *Salmonella typhimurium* + 3-NPA vs Ctrl+ *Salmonella typhimurium* **p=0.009. Cerebellum: Ctrl+ *Salmonella typhimurium* and mix Flavonoids+ *Salmonella typhimurium* vs mix Flavonoids+ *Salmonella typhimurium* +3-NPA **p=0.002. Striatum, Duodenum and Stomach p=N.S.

## DISCUSSION

Recent studies reveal that rutin suppresses the production of tumor necrosis factor-α (TNF-α) and inhibits the lipopolysaccharide (LPS)-induced activation of nuclear factor-κB (NFκB) [31]. Neuroinflammatory responses involve the activation of the interleukins. In this study, interleukin is decreased in the animals that received diosmine, hesperidin, rutin and *Salmonella typhimurium*. Hence, we suggest that the successful colonization of *Salmonella typhimurium may* enable the rational design of effective therapeutic strategies [32]. These findings were in line with the results obtained by Wu et al, [33], who in their study reported that rutin significantly reduced the levels of reactive oxygen species and improved locomotion recovery. They suggest that in appropriate dosage conditions, the mechanism may be related to the alleviation of inflammation and oxidative stress.

GABA levels increased in animals treated with *Salmonella typhimurium* alone or combined with flavonoids plus Rutin® in cerebellum regions. This result is in agreement with the findings of other authors who suggest that GABA could be a good strategy to modulate immunological response in various inflammatory diseases, produced by microbial strain [34].

Transmitter dopamine metabolism by monoamine oxidase enzyme has been attributed to striatal damage in Huntingtońs disease (HD) model associated with mitochondrial toxin [35]. HD is a destructive neurodegenerative disorder associated with progressive loss of neuronal functions, which eventually bring about the death of specific brain parts with the striatum and cerebral cortex being the principal targets [36]. These results can be in line with the reports of the present study, for the fact that dopamine levels diminished in cortex regions of animals that received 3-nitropropionic acid treatment in combination with *Salmonella typhimurium* ATCC14028.

With regard to the animals treated with diosmine, hesperidin, rutin and *Salmonella typhimurium*, ATPase activity decreased in cerebellum, probably as consequence of changes in the affinity of the enzyme [37]. In the animals that received the same treatment plus 3-NPA, there was a decrease in lipoperoxidation in cerebellum, and this may be due to the fact that reactive oxygen species is the primary event in 3-NPA toxicity [38]. GSH levels decreased in the duodenum of the animals treated with 3-NPA alone or combined. These results may have relation with the reports of Kumar et al. [39], who suggest that 3NPA depleted the GSH in cortex.

Ca²⁺/Mg²⁺-dependent ATPase increased in the cerebellum of animals that received 3-NPA alone or combined. This result is in line with the reports of Naziroğlu et al. [40], who suggest that increased Ca ^2+^-ATPase activities is due to substances that induced brain injury by exhibiting free radical production, regulating calcium-dependent processes and supporting the antioxidant redox system.

The Catalase activity decreased in the animals treated with diosmine, hesperidine and rutin in cortex and cerebellum regions, but increased in the presence of 3-NPA. These results coincide with the reports of Mascaraque et al. [41], who suggest that rutin has a significant protective effect.

Huntington’s disease is an inherited neurodegenerative disease. It is characterized by excessive motor movements couple with cognitive and emotional deficits [42]. In addition, there is a marked neuronal loss among the medium-sized projection neurons of the dorsal striatum. In this study, however; diosmine, hesperidine, rutin and *Salmonella typhimurium* supplemented *in vivo*, protected the striatum. This suggests a mechanism that involves antioxidant activity by controlling the expression of antioxidant enzymes and other chaperones regulating proteostasis, with potential neuroprotective role [43]. Besides, histological changes revealed marked lesions of neuronal cells in experimental animals treated with nitropropionic acid.

## CONCLUSION

The protective role of antioxidant compounds by inhibiting inflammatory response and correcting the fundamental oxidant/antioxidant imbalance in patients suffering from neurodegenerative diseases are important vistas for further research.

We recommend further studies to investigate the possible relationship between the flavonoids, rutin, inflammation by LPS and 3-NPA in different animal models. As a possible protective barrier against pro-inflammatory responses, it may be a new dietary strategy to combat Huntington’s disease.

Abbreviations.

Anova: Analysis of variance
CNS: Central nervous system
UFC: Colony-forming units
Rutin: Diosmine/hesperidine
DA: Dopamine
GABA: γ-Aminobutyric acid
GSH: Glutathione
5-HIAA: 5-Hydroxyindole acetic acid
3-NPA: 3-Nitropropionic acid
NOGSH: Nitroso-glutathione
RNS: Reactive nitrogen species
ROS: Reactive oxygen species
Na^+^, K^+^ ATPase: Sodium potassium ATPase
TBARS: Thiobarbaturic acid reactive substants

## DECLARATIONS

### Conflicts of interest/Competing interests

The authors declare they have no financial interests. The authors have no conflicts of interest relevant to the content of this article to declare.

### Funding

The authors did not receive support from any organization for the submitted work.

### Ethics

Animal management and caring was conducted according to the National and International guidelines of animal care. The protocol was approved by The Animal Committee of National Institute of Pediatrics with the reference number 026/2022.

### CRediT author statement

All the authors (David Calderón Guzmán, Norma Osnaya Brizuela, Maribel Ortiz Herrera, Hugo Juárez Olguín, Armando Valenzuela Peraza, Alberto Rojas Ochoa, Ernestina Hernández García, Rafael Coria Jiménez, Daniel Santamaria del Angel) participated in substantial manner in many areas of the present study beginning from its conception to drafting and critically reviewing the works, and to approving the final draft and signing for its publication as well as in choosing the journal in which the article has to be submitted.

### Accountability statement

All authors agreed to be answerable for the accuracy and integrity of the data, the originality of the work and proper attribution of all sources, the intellectual content and critical revisions of the manuscript, as well as adherence to reporting standards.

## Acknowledgments

We thank Dr. Cyril Ndidi Nwoye Nnamezie, an expert translator and a native English speaker, for his help in preparing this manuscript. Also, our thanks go to Instituto Nacional de Pediatria in facilitating all necessary avenues for the publication of this article.

## Availability of data and material

Any data and material used in this study are available on request to the correspondence author.

## Notes

### Competing Interest Statement

The authors have declared no competing interest.

## REFERENCES

1. Rami A, Ferger D, Krieglstein J. Blockade of calpain proteolytic activity rescues neurons from glutamate excitotoxicity. Neurosci Res 1997;27: 93–97.

2. Aliev G, Obrenovich ME, Tabrez S, et al. Link between cancer and Alzheimer disease via oxidative stress induced by nitric oxide-dependent mitochondrial DNA overproliferation and deletion. Oxid Med Cell Longev 2013; 962984. doi: 10.1155/2013/962984.

3. Tariq M, Khan HA, Elfaki I, Al Deeb S, Al Moutaery K. Neuroprotective effect of nicotine against 3-nitropropionic acid (3-NP)-induced experimental Huntington’s disease in rats. Brain Res Bull 2005;67(1-2): 161–8. doi: 10.1016/j.brainresbull.2005.06.024.

4. Guay DR. Tetrabenazine, a monoamine-depleting drug used in the treatment of hyperkinetic movement disorders. Am J Geriatr Pharmacother 2010;8(4): 331–73. doi: 10.1016/j.amjopharm.2010.08.006.

5. Hogg N, Singh RJ, Kalyanaraman B. The role of glutathione in the transport and catabolism of nitric oxide. FEBS Let 1996;382: 223–228.

6. Beckman JS, Beckman TW, Chen J, Marshall PA, Freeman BA. Apparent hydroxyl radical production by peroxynitrite: Implications for endothelial injury from nitric oxide and superoxides. Proc Natl Acad Sci USA 1990;87: 1624–1629.

7. Gutteridge JM, Halliwell B. The measurement and mechanism of lipid peroxidation in biological systems. Trends Biochem Sci 1990; 15:129–135

8. Driver AS, Kodavanti PR, Mundy WR. Age-related changes in reactive oxygen species production in rat brain homogenates. Neurotoxicol Teratol 2000;22: 175–181.

9. Vogt MC, Brüning JC. CNS insulin signaling in the control of energy homeostasis and glucose metabolism - from embryo to old age. Trends Endocrinol Metab 2012, [Epub ahead of print).

10. Guiney DG. The role of host cell death in Salmonella infections. Curr Top Microbiol Immunol 2005;289: 131–50. doi: 10.1007/3-540-27320-4_6.

11. Qiao L. Mechanisms for the Invasion and Dissemination of Salmonella. Can J Infect Dis Med Microbiol 2022; 2655801. doi: 10.1155/2022/2655801.

12. Ju Jung, Ryong Kim. Beneficial Effects of Flavonoids Against Parkinson’s Disease. J Med Food 2018;21(5): 421–432. doi: 10.1089/jmf.2017.4078.

13. Burda S, Oleszek W. Antioxidant and antiradical activities of flavonoids. J Agric Food Chem 2001;49: 2774–2779.

14. Cushnie TPT, Lamb AJ. Antimicrobial activity of flavonoids. Int J Antimicrob Agent 2005;26: 343–356.

15. Soukop J, Večeřa R. Selected polyphenolic compounds and their use as a supportive therapy in metabolic síndrome. Ceska Slov Farm 2022;71(4): 137–141

16. Bai M, Radhakrishnan A, Haleagrahara N. Rutin, a bioflavonoid antioxidant protects rat pheochromocytoma (PC-12) cells against 6-hydroxydopamine (6-OHDA)-induced neurotoxicity. Int J Mol Med 2013;32(1): 235–40. doi: 10.3892/ijmm.2013.1375.

17. Marafiga L, Lopes M, Franzen da Silva A, et al. Rutin protects Huntington’s disease through the insulin/IGF1 (IIS) signaling pathway and autophagy activity: Study in Caenorhabditis elegans model. Food Chem Toxicol 2020;141: 111323. doi: 10.1016/j.fct.2020.111323.

18. Swapna I, Sathya KV, Murthy CR, Senthilkumaran B. Membrane alterations and fluidity changes in cerebral cortex during ammonia intoxication. Neuro Toxicol 2005;335: 700–704.

19. Stefanello FM, Chiarani F, Kurek AG. Methionine alters Na^+^, K^+^ ATPase activity, lipid peroxidation and nonenzymatic antioxidant defenses in rat hippocampus. Int J Dev Neurosc 2005;23: 651–656.

20. Calderon GD, Juarez OH, Hernandez GE, et al. Effect of an antiviral and vitamins A,C,D on dopamine and some oxidative stress markers in rat brain exposed to ozone. Arch Biol Sci Belgrade 2013;65(4): 1371–1379.

21. Thygesen P, Brandt L, Jsrgensen T, et al. Immunity to experimental Salmonella typhimurium infections in rats. Transfer of immunity with primed CD4+CD2S^high^ and CD4+CD25 ^low^T lymphocytes. APMIS 1994;102: 489–494.

22. Calderón GD, Osnaya BN, García AR, Hernández GE, Guillé PA. Levels of glutathione and some biogenic amines in the human brain putamen after traumatic death. Proc West Pharmacol Soc 2008;51: 25–2.

23. Hsieh CY, Tsai EM, Wu HL. Simple and sensitive liquid chromatographic method with fluorimetric detection for the analysis of gamma-amino-n-butyric acid in human urine. Anal Chim Acta 2006;577: 201–206.

24. Guzman DC, Garcia EH, Brizuela NO, Jimenez FT. Effect of oseltamivir on catecholamines and some oxidative stress markers in the presence of oligoelements in rat brain. Arch Pharm Res 2010;33: 1671–1677.

25. Hissin PJ, Hilf R. A flurometric method for determination of oxidized and reduced glutathione in tissue. Anal Biochem 1974;4: 214–226.

26. Calderón-Guzmán D, Espitia-Vázquez I, Juárez-Olguín H, et al. Effect of toluene and nutritional status on serotonin, lipid peroxidation levels and Na^+^/K^+^ATPase in adult rat brain. Neurochem Res 2005;30: 619–624.

27. Fiske CH, Subbarow Y. The colorimetric determination of phosphorus. J Biol Chem 1925;66: 375–400.

28. Sinha AK. Colorimetric assay of catalase. Anal Biochem 1972;47: 389–394.

29. Luna LT. Manual of Histologic Staining Methods of the Armed Force Institute of Pathology. McGraw Hill Book Co., New York, 1968; pp: 1–3926.

30. Castilla-Serna L. Manual Práctico de Estadística para las Ciencias de la Salud. Editorial Trillas. 2011;1° Edición. México, D.F.

31. Lee W, Ku AK, Bae JS. Barrier protective effects of rutin in LPS-induced inflammation in vitro and in vivo. Food Chem Toxicol 2002;50(9): 3048–55. doi: 10.1016/j.fct.2012.06.013.

32. Zhang Z, Liu S, Huang J, et al. Phloretin is protective in a murine salmonella enterica serovar typhimurium infection model. Microb Pathog 2021:161(Pt B): 105298. doi: 10.1016/j.micpath.2021.105298.

33. Wu J, Maoqiang L, Fan H, et al. Rutin attenuates euroinflammation in spinal cord injury rats. J Surg Res 2016;203(2): 331–7. doi:10.1016/j.jss.2016.02.041.

34. Sokovic S, Djokic J, Dinic M, et al. GABA-Producing Natural Dairy Isolate From Artisanal Zlatar Cheese Attenuates Gut Inflammation and Strengthens Gut Epithelial Barrier in vitro. Front Microbiol 2019;10: 527. doi:10.3389/fmicb.2019.00527.

35. Smith RR, Dimayuga ER, Keller JN, Maragos WF. Enhanced toxicity to the catecholamine tyramine in polyglutamine transfected SH-SY5Y cells. Neurochem Res 2005;30(4): 527–31.

36. Browne SE, Beal MF. Oxidative damage in Huntington’s disease pathogenesis. Antioxid Redox Signal 2006;8(11-12): 2061–73.

37. Hoskins B, Ho IK, Meydrech EF. Effects of aging and morphine administration on calmodulin and calmodulin-regulated enzymes in striata of mice. J Neurochem 1985;44(4): 1069–73.

38. Mandavilli BS, Boldogh I, Van Houten B. 3-nitropropionic acid-induced hydrogen peroxide, mitochondrial DNA damage, and cell death are attenuated by Bcl-2 overexpression in PC12 cells. Brain Res Mol Brain Res 2005;133(2): 215–23.

39. Kumar P, Kalonia H, Kumar A. Protective effect of sesamol against 3-nitropropionic acid-induced cognitive dysfunction and altered glutathione redox balance in rats. Basic Clin Pharmacol Toxicol 2010;107(1): 577–82.

40. Naziroğlu M, Kutluhan S, Yilmaz M. Selenium and topiramate modulates brain microsomal oxidative stress values, Ca2+-ATPase activity, and EEG records in pentylentetrazol-induced seizures in rats. J Membr Biol 2008;225(1-3): 39–49.

41. Mascaraque C, Aranda C, Ocón B, et al. Rutin has intestinal antiinflammatory effects in the CD4+ CD62L+ T cell transfer model of colitis. Pharmacol Res 2014;90: 48–57. doi: 10.1016/j.phrs.2014.09.005.

42. Marafiga L, Valandro M, Franzen A, et al. Neuroprotective effects of rutin on ASH neurons in Caenorhabditis elegans model of Huntington’s disease. Nutr Neurosci 2022;25(11): 2288–2301. doi: 10.1080/1028415X.2021.1956254.

43. Ariano M, Wagle N, Grissell A. Neuronal vulnerability in mouse models of Huntington’s disease: membrane channel protein changes. J Neurosci Res 2005;80(5): 634–45. doi: 10.1002/jnr.20492.

